# XL-Ranker: A Computational Workflow for Prioritizing Protein-Protein Interactions from Cross-Linking Mass Spectrometry Data

**DOI:** 10.1101/2025.07.18.665625

**Authors:** Wenrong Chen, John M. Elizarraras, Fenglong Jiao, Lan Huang, Bing Zhang

## Abstract

Protein-protein interactions (PPIs) are central to virtually all biological processes, and their disruption can lead to a wide spectrum of human diseases. Cross-linking mass spectrometry (XL-MS) enables proteome-scale detection of interacting peptide pairs, from which PPIs can be inferred. However, interpretating XL-MS data to assign homologous cross-linked peptide pairs to specific protein interactions can be challenging. A major hurdle arises when a single peptide can map to multiple proteins, and when two such peptides are paired, the number of possible protein-protein interactions increases combinatorially. This mapping ambiguity can lead to inflated interaction networks that could compromise downstream analysis and biological interpretation. However, this problem has often been overlooked or addressed using heuristic approaches in previous XL-MS studies. To tackle this challenge, we developed XL-Ranker, a computational framework that combines a set cover graph algorithm and machine learning to systematically resolve peptide-mapping ambiguity and infer high-confidence PPIs from XL-MS data. Applied to an XL-MS dataset from HEK293 cells, XL-Ranker identified a high-confidence network with 880 PPIs involving 964 unique genes. Among all possible PPIs, the ones selected by XL-Ranker for inclusion in the final network had significantly higher interaction scores in the STRING database than the excluded ones. Network analysis further demonstrated that these interactions form biologically meaningful clusters, supporting the accuracy of our approach. In summary, XL-Ranker provides a practical solution to a key analytical challenge in XL-MS data interpretation, enhancing the reliability of PPI discovery.

## Introduction

Protein-protein interactions (PPIs) are fundamental to nearly all cellular processes, including signaling, gene regulation, metabolism, and structural organization^1,2^. Aberrant PPIs can disrupt essential biochemical pathways involved in cellular homeostasis, growth and proliferation, ultimately leading to diverse human diseases^3^. Therefore, accurate mapping of PPIs, particularly within living cells, is essential for understanding of the assembly, structure, functionality, and dynamics of protein networks, providing critical molecular insights linked to cell biology, human pathology, and therapeutic targeting.

Several methods are currently used to identify PPIs, including protein microarrays^4^, yeast two-hybrid assays^5,6^, affinity purification mass spectrometry (AP-MS)^7–10^, and cross-linking mass spectrometry (XL-MS)^11^. Among these, XL-MS has emerged as a powerful approach due to its high sensitivity, accuracy and unique ability to generate detailed, in vivo network topology maps of PPIs^11,12^. In particular, XL-MS is uniquely capable of detecting weak or transient interactions and those involving proteins that are difficult to solubilize, owing to its chemical crosslinking-based methodology^12–17^. Recent advancements in mass spectrometry (MS) instrumentation and computational algorithms^18–23^ for identifying cross-linked peptides have further enhanced the resolution and accuracy of protein complex topology mapping.

Despite the promise of XL-MS technology for large-scale interaction mapping, data interpretation remains a significant challenge^24,25^. One major obstacle is the ambiguity in assigning cross-linked peptide pairs to definitive PPIs, when peptide sequences are located at homologous regions shared by multiple proteins. Specifically, a single peptide can map to multiple proteins, and when two such peptides are cross-linked, the number of possible protein–protein interactions increases combinatorially (**Figure 1A**). This mapping ambiguity can lead to inflated interaction networks, which compromise downstream analyses and biological interpretation.

**Figure 1.**
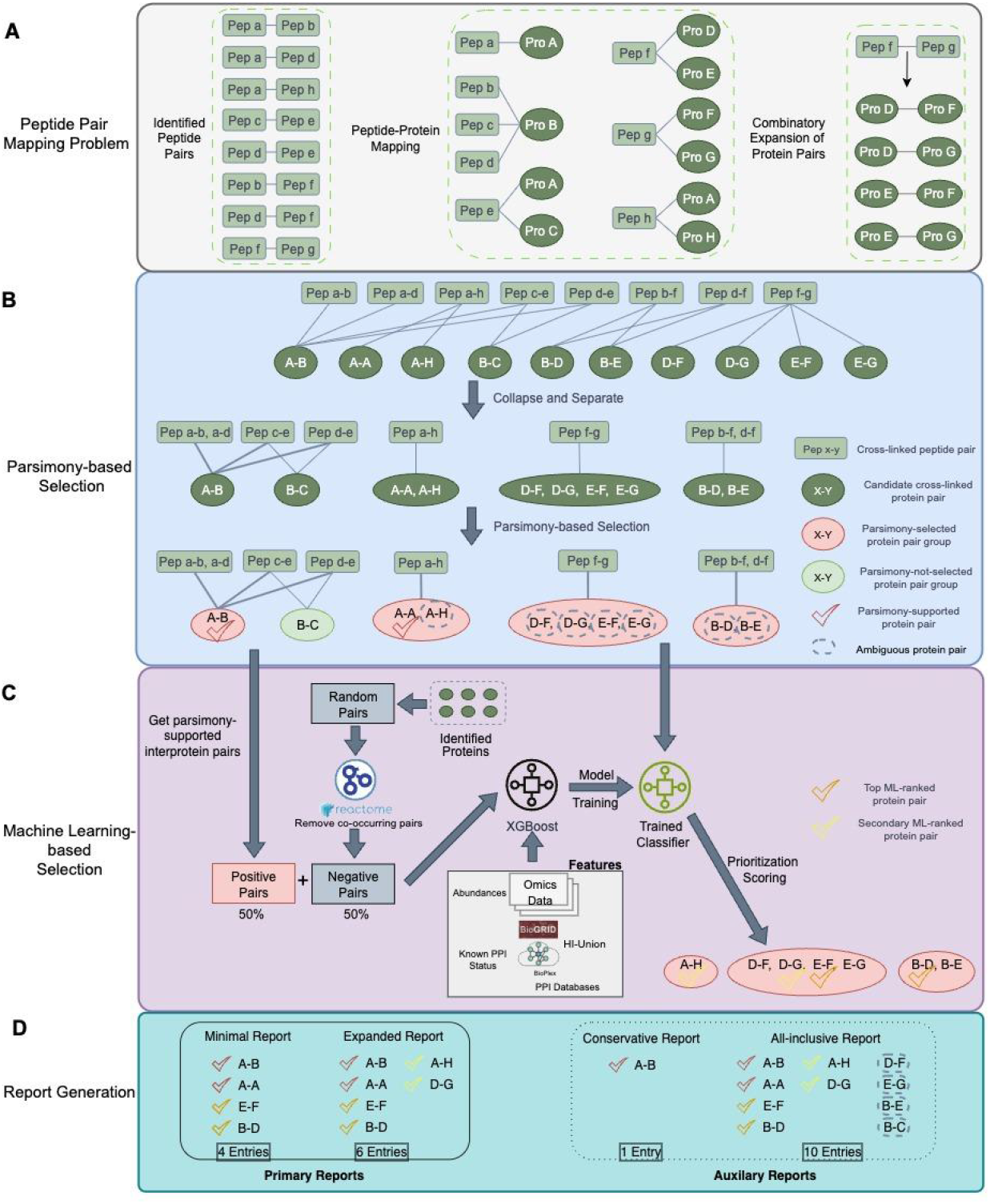
Systematic overview of the XL-Ranker workflow for PPI prioritization and selection from cross-linked peptides. **(A)** Peptide Pair Mapping Problem. Identified cross-linked peptide pairs are mapped to their corresponding proteins. Due to shared peptide sequences, resulting in combinatorial explosion of possible protein pairs. **(B) (C)** denotes the two panels of XL-Ranker workflow. **(B)** Parsimony-based selection panel. Identified peptide pairs and protein pairs are represented as a bipartite graph, with edges indicating sequence matches. Protein and peptide nodes with identical connection patterns are grouped together (Collapse step). The graph is then divided into connected components (Separate Step). Within each component, a minimal set of protein pair groups is selected to represent all observed peptide pairs (Parsimony-based selection step). In distinguishable protein pair groups, parsimony-selected protein pair is colored red and parsimony-not-selected pair is colored green. **(C)** ML-based selection panel. Indistinguishable protein pairs are further prioritized using a machine learning-based approach. The positive set for the model is the protein pairs selected during parsimony selection, while negative set contains random pairs excluding those sharing same reactome pathway or Gene Ontology cellular-component annotations. Positive and negative pairs are used to train an XGBoost classifier using features from abundance data and known PPI status from databases including BioGRID and Hi-Union. The model assigns prioritization scores to each protein pair and the highest scoring pair from each group is selected to construct the final PPI network. The final PPI network includes both parsimony-selected and ML-selected pairs. **(D)** Report generation panel. There are two primary reports including minimal report and expanded report generated by XL-Ranker. The two auxiliary reports including conservative report and all-inclusive report are provided for users as well.

Although widely recognized, this challenge remains inadequately addressed in previous studies. There is no standardized approach for resolving ambiguity when aggregating cross-linking data from residue-pair or peptide-pair level to PPI level. Methods used across published studies and computational tools are often undocumented or poorly described. At a high level, common approaches include reporting all possible protein pairs, restricting to unambiguous pairs only, or selecting a representative protein pair for each peptide pair—often based on random choice or alphabetical order of protein/gene names.

Here, we present XL-Ranker, a standardized computational workflow designed to systematically prioritize and select PPIs from cross-linked peptides. XL-Ranker consists of two steps: First, a parsimony-based selection strategy inspired by Occam’s razor is used to identify a minimal set of protein pairs needed to explain all observed cross-linked peptide pairs. This approach simplifies the interaction space while preserving explanatory power. Second, the remaining ambiguous protein pairs are further scored using an XGBoost machine learning model, which incorporates multiple discriminative features to assign confidence scores for candidate PPIs. XL-Ranker provides multiple reporting schemes with varying stringency levels to suit different applications. We applied XL-Ranker to XL-MS data from HEK293 cells and showed that its two-step prioritization strategy identifies high-confidence PPIs that assemble into biologically coherent modules, demonstrating its effectiveness in resolving peptide-to-protein ambiguity.

## Results and Discussion

### Overview of XL-Ranker workflow

XL-Ranker employs a two-stage computational framework to prioritize PPIs from cross-linked peptide data (**Figure 1**), aiming to systematically resolve peptide-to-protein ambiguity and generate interaction networks with varying levels of stringency.

In the first stage (**Figure 1B**), cross-linked peptides and their corresponding protein pairs are represented as a bipartite graph, with edges connecting peptide pairs to protein pairs they map to. Nodes with identical connection patterns are grouped, and the graph is decomposed into independent components. Within each component, a parsimony-based selection strategy is used to identify the minimal set of protein pair groups required to explain all observed peptide pair groups. If a protein pair group contains only a single protein pair, it is directly selected, as it unambiguously explains the corresponding peptide pair. In protein pair groups with multiple candidates, intraprotein pairs are selected when present for simplicity, as they introduce fewer nodes into the final network. We define protein pairs emerging from this step as parsimony-supported protein pairs.

The remaining ambiguous protein pairs are passed to the second stage (**Figure 1C**), where an XGBoost classifier is applied to further prioritize candidates. The classifier is trained using parsimony-supported interprotein pairs as positive examples and randomly sampled protein pairs—drawn from the full list of proteins identified in the XL-MS data—as negatives. Discriminative features include RNA or protein abundance of the constituent proteins—derived from the same or related samples—and evidence of physical interactions in public databases. For each ambiguous peptide pair, the model outputs a top ML-ranked protein pair, along with additional secondary ML-ranked pairs that exceed a user-defined prediction score threshold.

Based on these results, XL-Ranker generates PPI reports at two primary levels of stringency (**Figure 1D**). The *Minimal report* includes a single protein pair for each peptide pair, selected from either parsimony-supported or top-ranked ML-scored candidates. The *Expanded report* builds upon the Minimal report by additionally including other protein pairs that meet a user-defined ML-score threshold. In addition, XL-Ranker provides two auxiliary reports: a *Conservative report*, which includes only protein pairs unambiguously supported by the identified peptide pairs, and an *All-inclusive report*, which lists all possible protein pairs associated with each peptide pair.

### Application of XL-Ranker to XL-MS data from HEK293 cell line

To assess the performance of XL-Ranker, we applied it to crosslinked peptides identified by XL-MS analysis of the HEK293 cell line, including a total of 8,068 distinct peptide pairs.

These peptide pairs were mapped to the human Swiss-Prot protein database (version 2019.04), resulting in 8,779 candidate protein pairs, comprising 6,844 intraprotein interactions and 1,935 interprotein interactions.

Parsimony-based analysis of the corresponding bipartite graph prioritized 2,351 protein pair groups, from which 2,238 parsimony-supported protein pairs were selected, including 1,794 intraprotein pairs and 444 interprotein pairs. The remaining 2,076 ambiguous protein pairs were further evaluated using the ML model, using the protein abundance and known PPI status as input features. This analysis identified 156 top ML-ranked protein pairs, along with 280 secondary ML-ranked pairs exceeding a prediction score threshold of 0.5.

Based on these results, XL-Ranker generated four PPI reports with markedly different protein and interaction counts (**Table 1**). The Expanded report, which balances coverage and reliability, included 2,164 proteins and 2,674 interactions, comprising 880 interprotein interactions and 1,794 intraprotein interactions. Corresponding network of interprotein interactions is visualized in **Figure S1** as the XL-PPI network.

**Table 1.** Summary of four reports generated by XL-Ranker using HEK293 cell line data set.

| Report | Nodes | Edges | Intraprotein Edges | Interprotein Edges |
| --- | --- | --- | --- | --- |
| Minimal | 1992 | 2351 | 1751 | 600 |
| Expanded | 2164 | 2674 | 1794 | 880 |
| Conservative | 1785 | 2033 | 1589 | 444 |
| All-inclusive | 4,753 | 8,779 | 6,884 | 1,935 |

To evaluate the performance of the two main steps of XL-Ranker, we used the STRING scores to compare interprotein pairs selected and not selected during the parsimony-based and ML-based selection steps. Protein pairs selected in the parsimony step had a median STRING score above 0.95, while those not selected had a median score below 0.4 (**Figure 2B**). Similarly, protein pairs retained in the ML step had a median STRING score above 0.9, compared to a median of approximately 0.25 for those not-selected (**Figure 2B**). Overall, protein pair classification by XL-Ranker was well separated by STRING scores (**Figure 2C**), supporting the biological relevance of the selection criteria.

**Figure 2.**
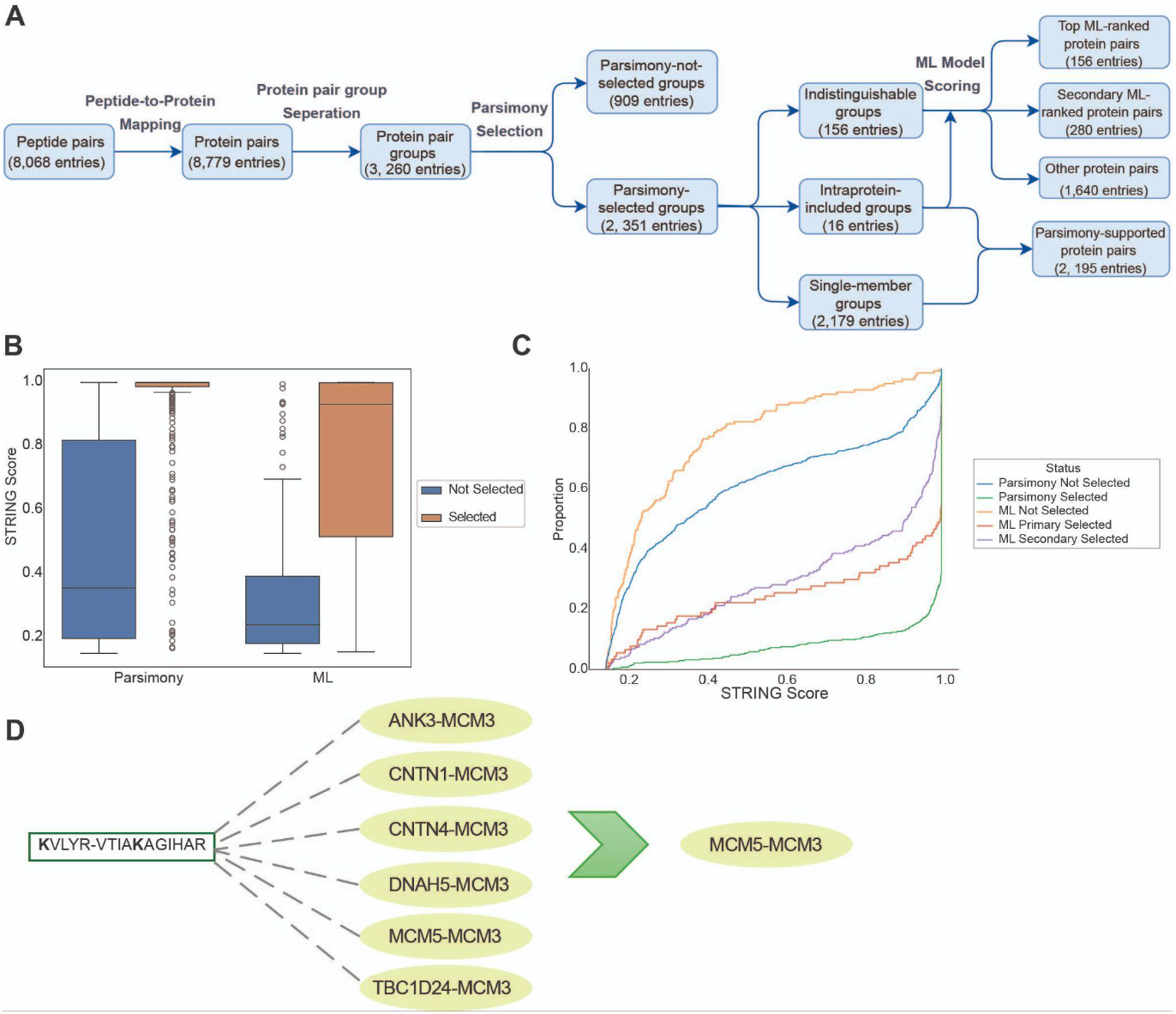
Evaluation of XL-Ranker using cross-linked peptides generated from the HEK293 cell line. **(A)** Number of interactions at each step of XL-Ranker workflow. **(B)** Distribution of STRING score differences between the selected and not-selected protein pairs in the parsimony and ML steps. **(C)** Comparison of STRING scores between parsimony-selected pairs vs not-selected pairs, and ML-selected pairs vs not-selected pairs. **(D)** The example for cross-linked peptides KVLYR-VTIAKAGIHAR. There are six possible PPIs mapped to this peptide pair and MCM5-MCM3 was prioritized by XL-Ranker.

XL-Ranker effectively reduced PPI ambiguity in the HEK293 XL-MS data set. For example, the peptide linkage KVLYR-VTIAKAGIHAR was initially associated with six possible PPIs. XL-Ranker prioritized the interaction between MCM3 and MCM5 (**Figure 2D**). These two proteins co-localize in the nuclear chromosome compartment and are known to interact functionally in processes such as DNA replication and transcriptional regulation^26^. In contrast, the alternative candidates (ANK3, CNTN1, CNTN4, DNAH5 and TBC1D24) do not share the same subcellular localization with MCM3, making their potential interaction with MCM3 highly unlikely.

### Functional analysis of the PPI network

To further investigate the functional relevance of the XL-PPI network, we applied the NetSAM algorithm to uncover its hierarchical organization. We identified 4 hierarchical levels and 71 modules with at least 5 genes. Among these modules, 55 significantly overlapped with at least one GO annotation (FDR < 0.05, Fisher’s exact test followed by Benjamini-Hochberg adjustment), indicating their functional coherence. We categorized those GO annotations to 7 functional categories including metabolism, development & differentiation, immunity, membrane transport, gene regulation, cytoskeleton & cell structure and protein homeostasis (**Figure 3A**).

**Figure 3.**
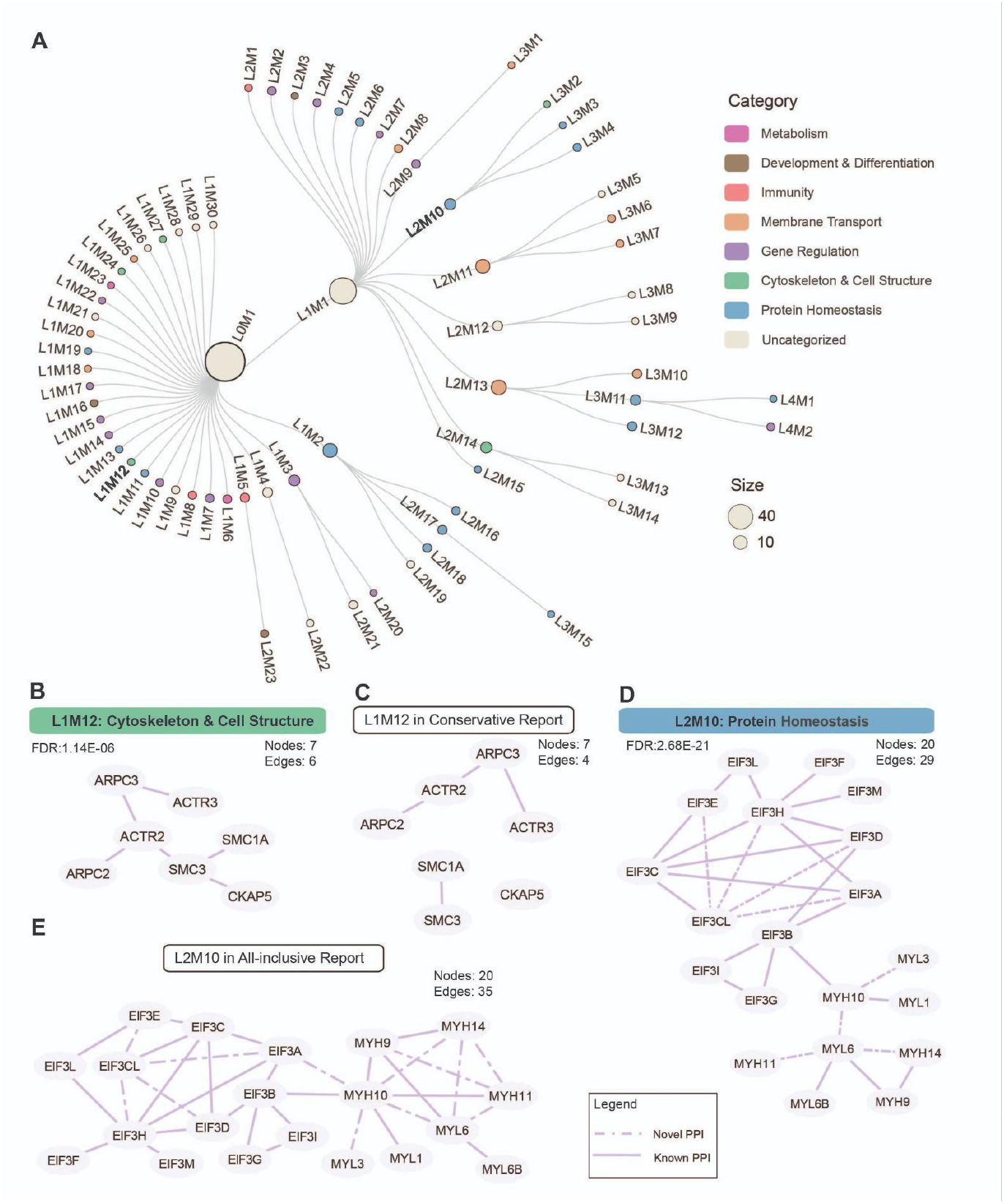
Functional analysis based on hierarchical modular structure of PPI network constructed with XL-Ranker. **(A)** Hierarchical modular structure. Nodes represent NetSAM-derived modules with node size proportional to the number of genes in the module. Nodes are colored using different categories from functional enrichment results on the basis of genes in the module. **(B)(C)** Node-link diagram of module L1M12 in expanded report and conservative report. **(D)(E)** Node-link diagram of L2M10 in expanded report and all-inclusive report. The solid edges represents known PPI and dash-dot egdes represents novel PPI.

For instance, L1M12 is a cytoskeleton and cell structure module comprising six PPIs among seven genes: ACTR2, ACTR3, ARPC2, ARPC3, SMC1A, SMC3 and CKAP5 (**Figure 3B**). ACTR2, ACTR3, ARPC2 and ARPC3 form the ARP2/3 complex, which promotes branched actin filaments and is essential for maintaining cell shape and movement^27^. SMC1A and SMC3 are involved in chromosome cohesion but also affect cytoskeleton function^28^. All PPIs in the module are supported by previously reported physical interactions. For comparison, restricting to include PPIs derived from uniquely mapped cross-linked peptides (conservative report) reduced the module size to four PPIs among the seven genes within the module L1M12, resulting in the loss of two known PPIs (**Figure 3C**). This highlights the ability of XL-Ranker to balance coverage and reliability.

As another example, module L2M10, associated with protein homeostasis, comprises 29 PPIs involving 20 genes (**Figure 3D**). Of these, 21 PPIs (72.4%) are supported by previous publications. While the All-inclusive report included six additional edges for genes in L2M10, only two of them (33.3%) are supported by existing literature (**Figure 3E**). Thus, XL-Ranker reduced complex subnetworks to core interactions that better align with existing literature.

## Conclusions and Discussion

Accurate interpretation of cross-linked peptides from XL-MS data remains a critical challenge, largely due to the inherent ambiguity in mapping cross-linked peptides to their corresponding PPIs. In this study, we proposed XL-Ranker, a computational workflow designed to systematically prioritize the PPIs with ambiguity. By applying XL-Ranker to a XL-MS dataset from HEK 293 cell line, we demonstrated its ability to extract a high-confidence interaction network consisting of 880 interprotein interactions across 964 genes. Comparative analysis showed that the PPIs selected by XL-Ranker had significantly higher STRING scores than the excluded ones, indicating strong support from independent databases. Furthermore, network analysis revealed that the selected interactions formed coherent functional modules, aligning with known biological processes such as cytoskeleton and cell structure, gene regulation and protein homeostasis. These results validated the effectiveness of our method in resolving ambiguous mapping and improving the biological interpretation of XL-MS-derived PPI networks.

Despite its strength, several limitations should be noted. First, the quality of the constructed network depends on the quality of cross-linked peptide identifications and input abundance features. Inaccurate or low-confidence cross-linked peptide spectrum matches (CSM) may propagate errors through both selection steps. Second, the ML model uses external databases such as BioGRID, BioPlex and HI-Union to define training sets and features. This may introduce biases toward well-characterized interactions and limit discovery in less studied proteins or complexes.

In conclusion, XL-Ranker provides a systematical and reproducible solution to the key challenge in XL-MS data analysis. By reducing the mapping ambiguity and prioritizing biologically associated interactions, it improves the accuracy and utility of XL-PPI networks. XL-Ranker can serve as a valuable tool for the proteomics community, enabling more reliable discovery of PPIs and deeper insights into biological processes and therapeutic targets.

## Methods

### Data set preparation

The cross-linked peptides identified from the HEK293 XL-MS data set were acquired from a published study^29^. The RAW data of the global proteomic data set for HEK293 cell line was downloaded from PRIDE with PXD001468^30^. The protein database used in the database search was SwissProt-Human (20,421 entries, version 2019.04) which was harmonized with the XL-MS data processing. The MS/MS spectra were processed with FragPipe (v22.0) and the database search was performed using MS-Fragger (version 4.1) with default setting. The default search parameters are set as follows: set precursor-ion mass tolerance as 20 ppm, fragment mass tolerance as 20 ppm, fixed modifications as carbamidomethylation (+57.0215), variable modifications as methionine oxidation (+15.9949), N-terminal protein acetylation (+42.0106), allow for up to two missed cleavage sites.

### Parsimony-based Selection

Parsimony-based selection was used to reduce redundancy in peptide pair to protein pair mapping which is similar with the parsimony strategy applied on the peptide to protein mapping ^31^. All possible mapping between protein pairs and peptide pairs identified from the XL-MS data were organized into a bipartite graph consisting of two types of nodes: protein pair nodes and peptide pair nodes. Edges connected peptide pair nodes to protein pair nodes if the peptide pair sequences matched the protein pair sequences. Due to ambiguities in mapping peptides to proteins, connections within the graph could be complex. To simplify this bipartite graph, the protein pair nodes that shared identical set of connected peptide pair vertices were grouped together, as these protein pairs could not be differentiated based on peptide identification alone. Similarly, peptide pair nodes which connected to the same set of protein pair nodes were clustered into groups.

The simplified graph was further broken down into independent subgraphs by identifying connected components which were the largest subgraphs with interconnected nodes. Within each connected component, a minimal selection of protein pair groups was chosen through a greedy set cover algorithm to represent all associated peptide pair nodes in the groups as the parsimonious protein pair list. The selected protein pair groups containing a single pair resulted in that pair being parsimoniously selected. For multi-membered selected protein pair groups, intra pairs were prioritized, as they introduce fewer nodes into the final network. For protein groups containing multiple intra pairs, the protein with the highest abundance is selected, and for ties, the protein is selected alphabetically. The remaining inter protein pairs in the group are scored by the machine learning model for selection. For groups containing only inter protein pairs, all pairs were given to the machine learning model for selection.

### Machine learning (ML) model-based Selection

To address ambiguity that can not be resolved by parsimony selection, we train a XGBoost classifier to provide prioritization scores for the unresolved protein pairs. The positive dataset for the model are the interprotein pairs selected during parsimony selection. To create a negative set, we generate random protein pairs from the list of all possible proteins identified in the cross-linking data. Random pairs that contain proteins that occur in the same reactome pathway or Gene Ontology cellular-component are removed from the negative set ^32,33^. The model is trained with abundance data and known PPI status. Users can provide multiple omics data sets, typically RNAseq and proteomics data. For each input dataset, we use the abundance of both proteins in a pair as input features for the model. Additionally, we add a binary feature of if the pair was previously found to have physical interaction in the BioGRID, BioPlex or HI-Union databases ^9,34,35^.

XL-Ranker uses an ensemble of ten XGBoost classification models to provide more robust prediction scores. For a single model, training is performed using 5-fold cross-validation, with an equal number of positive and negative pairs in both the training and testing data. The negative pairs are randomly sampled from the larger pool of candidate negatives, with each model having a different set of negatives. Each model will also predict a score for each unresolved protein pair. After running the models, the final score for an unresolved pair is the mean across the 10 models. Protein pairs with a prediction score greater than 0.5 are selected by the ML model.

### Hierarchical module identification and functional enrichment analysis

The hierarchical module was identified using the NetSAM algorithm implemented in R (https://bioconductor.org/packages/release/bioc/html/NetSAM.html). The main function of the package takes a network edge list as input and outputs an ‘nsm’ file that contains all detected modules arranged hierarchically. Two key parameters are minModule and modularityThr. The minModule parameter controls the smallest allowed module size, defined as a fraction of the total number of nodes in the network. Modules smaller than this threshold are not further divided. In our analysis, we set minModule so that the minimum module size was 5 nodes with a modularity threshold of 0.2.

For functional enrichment, we used WebGestalt to perform over-representation analysis to identify the most significantly enriched Gene Ontology Biological Process for each hierarchical module. For modules with no gene set with an FDR < 0.05, no enriched set was reported^32,36^.

## Acknowledgements

This study was supported by the National Cancer Institute (NCI) CPTAC award U24 CA271076 (B.Z.) funding from the McNair Medical Institute at The Robert and Janice McNair Foundation (B.Z.) and NIH R35GM145249 to L.H.

## Code Availability

The source code of XL-Ranker can be found at GitHub: github.com/bzhanglab/xlranker

## Figures

**Figure S1.**
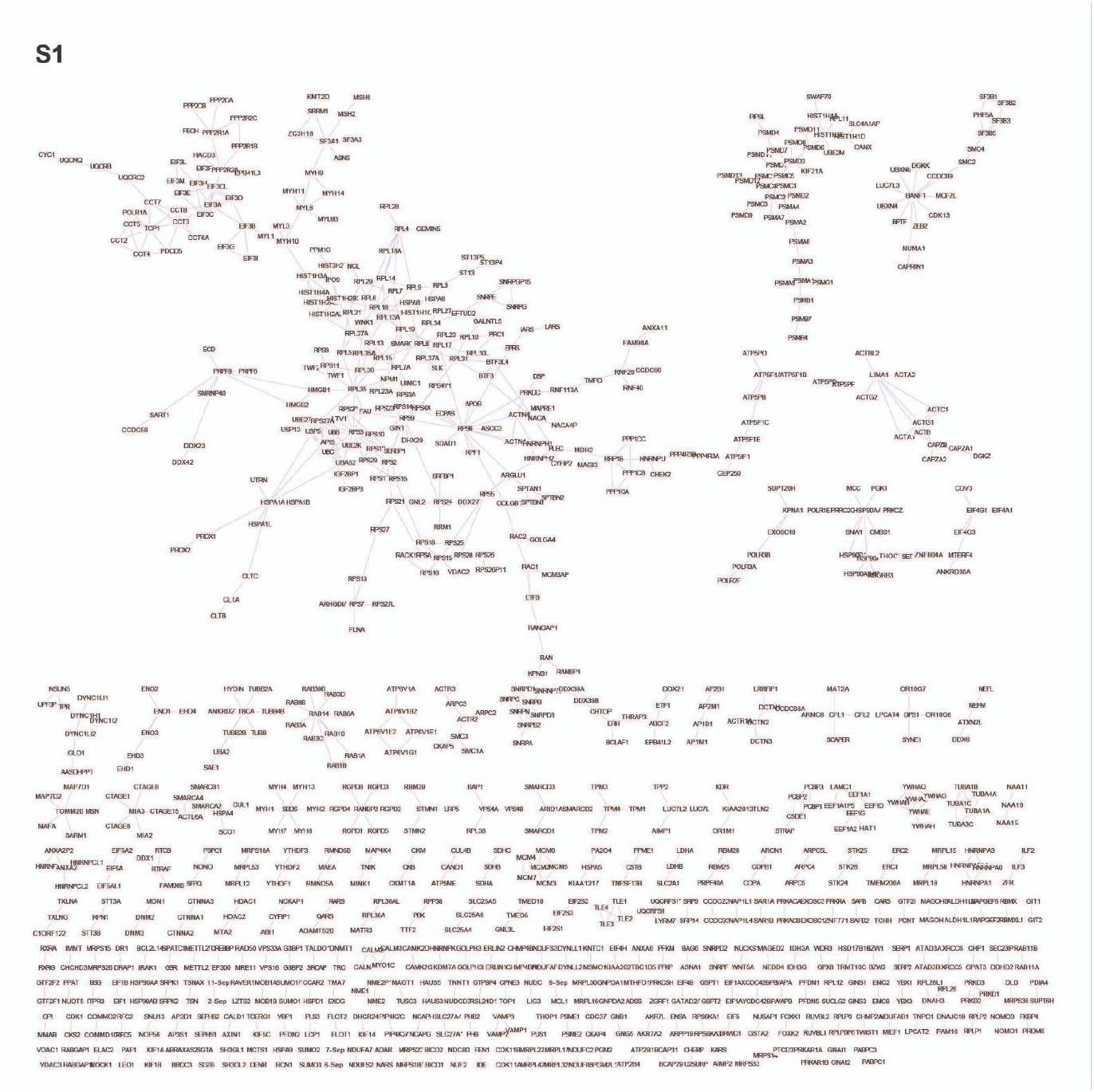
Final PPI network with interprotein interactions of HEK293 cell line. The network is composed of 964 nodes and 880 edges.

